# *Comamonas aquatica* inhibits TIR-1/SARM1 induced axon degeneration

**DOI:** 10.1101/2024.11.20.622298

**Authors:** Lauren C O’Connor, Woo Kyu Kang, Paula Vo, Jessica B Spinelli, Mark J Alkema, Alexandra B Byrne

**Affiliations:** Department of Neurobiology, University of Massachusetts Chan Medical School, Worcester, MA, 01605, USA; NeuroNexus Institute, University of Massachusetts Chan Medical School, Worcester, MA, 01605, USA; Program in Molecular Medicine, University of Massachusetts Chan Medical School, Worcester, MA, 01605, USA

**Keywords:** *C. elegans*, microbiome, gut-brain axis, neurodegeneration, TIR-1/SARM1, homocysteine, *Comamonas aq*

## Abstract

Emerging evidence suggests the microbiome critically influences the onset and progression of neurodegenerative diseases; however, the identity of neuroprotective bacteria and the molecular mechanisms that respond within the host remain largely unknown. We took advantage of *Caenorhabditis elegans’* well characterized nervous system and ability to eat uni-bacterial diets to determine how metabolites and neuroprotective molecules from single species of bacteria suppress degeneration of motor neurons. We found *Comamonas aquatica* significantly protects against degeneration induced by overexpressing a key regulator of axon degeneration, TIR-1/SARM1. Genetic analyses and metabolomics reveal *Comamonas* protects against neurodegeneration by providing sufficient Vitamin B12 to activate METR-1/MTR methionine synthase in the intestine, which then lowers toxic levels of homocysteine in TIR-1-expressing animals. Defining a molecular pathway between *Comamonas* and neurodegeneration adds significantly to our understanding of gut-brain interactions and, given the prominent role of homocysteine in neurodegenerative disorders, reveals how such a bacterium could protect against disease.

## Introduction

Changes in gut microbiota have been linked to neurodegenerative diseases and neurological conditions, including Alzheimer’s disease, Parkinson’s disease, Amyotrophic lateral sclerosis (ALS), Huntington’s disease, autism, anxiety, depression, addiction, schizophrenia, and migraine^1–7^. However, the molecular mechanisms that mediate gut-brain interactions have been exceedingly difficult to unravel, primarily due to the complexity of the mammalian nervous system and microbiome. Identifying microbiota that regulate neurodegeneration, as well as determining which neuronal mechanisms are directed by specific microbial metabolites, is crucial to understand how to manipulate the microbiome and target specific molecules to counteract debilitating neuronal disorders.

*C. elegans* and their bacterial diet are a potent and highly informative model for studying conserved host-microbiome interactions, as both are well defined, genetically tractable, and relatively easy to manipulate^8^. In addition, molecular mechanisms that regulate neuronal development and function are significantly conserved between *C. elegans* and multiple species. The *C. elegans* proteome is 83% conserved with humans, and consequently, up to 75% of human disease-related genes, including genes associated with Alzheimer’s disease, have homologs in *C. elegans*^9–11^. *C. elegans* encounter and feed on a variety of bacteria in nature; however, *C. elegans* can also be raised on uni-bacterial diets in the laboratory^12^. The ability to examine the consequences of specific naturally occurring bacteria on conserved animal physiology is a powerful approach to gain mechanistic insight into how particular diets influence development as well as reveal conserved mechanisms of disease^8,13–16^.

Neurodegenerative diseases and neurologic injury both lead to a devastating loss of cognitive or motor function caused by axon degeneration and consequent neuronal cell death. Age is a major risk factor for neurodegenerative diseases such as Alzheimer’s disease, Amyotrophic lateral sclerosis and Parkinson’s disease, among many others^17^. Our current understanding is that neurodegenerative conditions such as Alzheimer’s disease and neuropathies are thought to be “dying back” diseases where the portion of axon distal to the cell body is most susceptible to damage, which ultimately progresses back to the cell body^18^. Injury-induced degeneration recapitulates this dying-back phenotype. Wasting of the distal portion of an injured neuron was once considered to be a passive process, in which a lack of signaling and nutrients from the nucleus results in degeneration^19^. However, discovery of a mutant mouse in which degeneration is significantly delayed, called Wallerian degeneration slow (Wld^S^), indicated that degeneration is a molecular process^19,20^. The subsequent discovery of the prodegenerative gene *dSarm* in *Drosophila melanogaster* and its homolog *Sarm1* in mammals cemented the conclusion that Wallerian degeneration is actively regulated^21,22^.

We recently found that the *C. elegans* homolog of SARM1, TIR-1, also promotes degeneration of injured axons^23^. Overexpression of an active isoform of TIR-1/SARM1 (hereafter referred to as TIR-1(oe)), which lacks its N-terminal autoinhibitory domain, in GABAergic motor neurons causes spontaneous and significant motor axon degeneration^23^. Like axons undergoing Wallerian degeneration, TIR-1(oe) axons display thinning, beading, fragmentation and ultimately degenerate. Importantly, as with dSarm/SARM1, we found that TIR-1 requires its NADase activity to induce chronic degeneration, indicating *C. elegans* provides a novel and unexplored model with which to investigate conserved mechanisms of TIR-1/dSARM/SARM1 mediated degeneration^21–24^. A better understanding of how to disrupt constitutive TIR-1/SARM1 signaling is critical to designing effective clinical therapies for degenerative disease, as recent evidence demonstrates that mutations that disrupt the N-terminal autoinhibitory domain of human SARM1 also confer constitutive SARM1 NADase hyperactivity and contribute to Amyotrophic lateral sclerosis, Parkinson’s disease, negative outcomes of traumatic brain injury and Alzheimer’s disease^25–31^. While diet and the microbiome are thought to affect the onset and progression of these SARM1 dependent neurodegenerative conditions, very few precise protective or prodegenerative molecular interactions between the microbiome and the nervous system have been identified^32–34^.

We took advantage of the degenerative phenotype of TIR-1/SARM1 overexpressing animals to screen for bacterial diets that inhibit degeneration. We found that a diet consisting solely of *Comamonas aquatica* DA1877 (*Comamonas*) protects axons from chronic neurodegeneration induced by activated TIR-1/SARM1. *Comamonas* requires its ability to produce B12 to protect against TIR-1/SARM1 induced neurodegeneration. We found that B12 confers neuroprotection by interacting with the neuron-extrinsic function of the conserved methionine synthase enzyme METR-1/MTR. Liquid chromatography-mass spectrometry (LC-MS) analyses revealed a dramatic decrease in homocysteine levels in TIR-1(oe) animals raised on a *Comamonas* diet compared to animals raised on a standard *E. coli* OP50. Our results reveal a mechanism by which a single bacterium is able to confer neuroprotection by lowering toxic homocysteine levels, which are tightly associated with neurodegenerative disease.

## Results

To investigate causal interactions between bacterial diets and molecular mechanisms that regulate neurodegeneration, we screened non-pathogenic bacterial diets that *C. elegans* encounters in the wild for their ability to suppress motor axon degeneration (**Figure 1**). Wild-type animals raised on a standard laboratory *E. coli* OP50 diet have 16 GABA motor axons that extend circumferentially on their right side. These neurons develop in a stereotypical pattern that is readily observed *in vivo* using a GABA specific fluorescent reporter (*Punc-47::GFP*) (**Figure 1C, D**). We previously found that GABA axons degenerate in animals that express active TIR-1/SARM1 in their GABA motor neurons (hereafter referred to as TIR-1(oe)). Mature TIR-1(oe) animals have an average of only two intact axons when raised on the standard *E. coli* OP50 diet (p≤0.0001 relative to wildtype, Student’s t-test) (**Figure 1C, D)**^23^. Overexpression of SARM1, the human TIR-1 orthologue, lacking its autoinhibitory domain in *C. elegans* motor neurons also induces significant axon degeneration (**Figure 1A, Supplementary Figure 1**). Together with our previous finding that axon degeneration in TIR-1(oe) animals requires conserved amino acid residues E642 and E788, which are essential for the NADase activity of dSarm/SARM1^23^, our data suggest that overexpressing TIR-1 and SARM1 induces degeneration using conserved mechanisms. We screened bacterial diets for their ability to confer protection by quantifying the number of intact axons in mature TIR-1(oe) animals fed each individual diet from birth (**Figure 1D**). We discovered that a diet consisting solely of *Comamonas aquatica* (hereafter referred to as *Comamonas*) protects significantly more axons from TIR-1(oe) induced degeneration than the *E. coli* OP50 diet **(Figure 1D)**. These data reveal a *Comamonas* diet protects against motor axon degeneration that is induced by a key and conserved promoter of degeneration, TIR-1/SARM.

**Figure 1:**
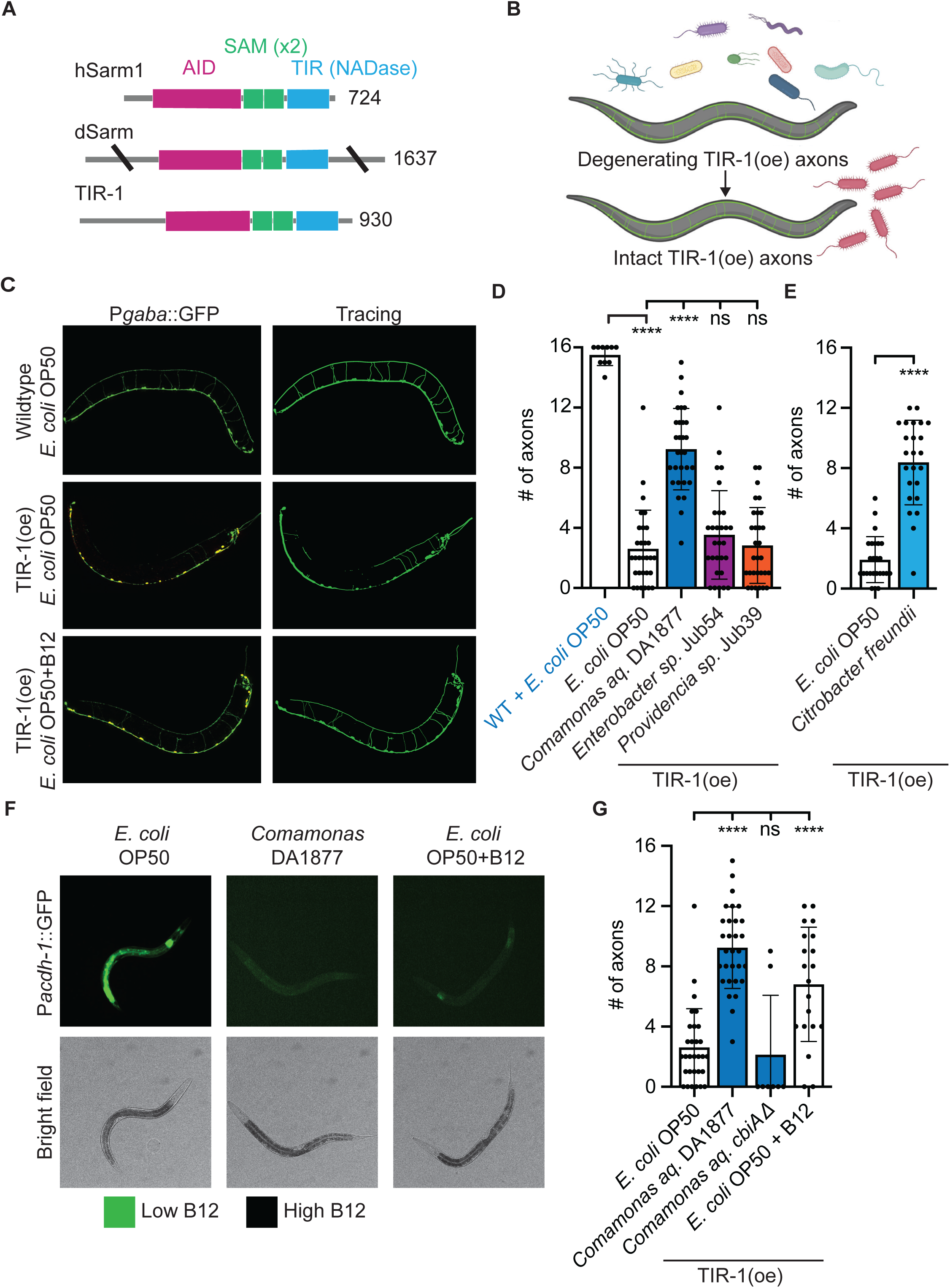
*Comamonas aquatica* protects against TIR-1 induced neurodegeneration. (**A**) *C. elegans* TIR-1 is a prodegenerative gene highly conserved with its orthologs hSarm1, and dSarm in humans and Drosophila. TIR-1 is comprised of four primary domains: an N-terminal autoinhibitory domain (AID), two SAM multimerization domains, and a TIR (toll-interleukin receptor) domain with NADase function. **(B)** *C. elegans* are capable of surviving on uni-bacteral diets, enabling direct investigation of whether and how specific bacterial diets protect against neurodegeneration. (**C**) Representative images of wild type and TIR-1(oe) animals fed standard *E. coli* OP50 diet. Wild-type animals have 16 GABA commissures on their right side. Overexpression of the active TIR-1b isoform, which lacks the autoinhibitory domain, in GABA motor axons induces significant axon degeneration. TIR-1(oe) animals fed B12 supplemented diet induces significant axon protection. (**D**) A screen of individual diets revealed that *Comamonas aq.* conferred significant protection against TIR-1 mediated degeneration (N=10, 30, 30, 30, 30). (**E**) Microbe *Citrobacter freundii* isolated from the human microbiome, exerts neuroprotection on TIR-1 mediated degeneration of the GABAergic motor neurons (N=25,24). (**F**) Representative images of animals expressing the B12 sensor P*acdh-1*::GFP, which reflects the amount of B12 absorbed from the diet (GFP expression=low B12). Both a *Comamonas aq.* diet and B12 supplementation to OP50 plates provide sufficient amounts of B12 to suppress the P*acdh-1*::GFP sensor in wild-type animals, indicating they are capable of absorbing B12. **(G)** A mutant *Comamonas aq.* strain that is unable to produce B12 (*Comamonas aq. cbiA*Δ) does not protect against axon degeneration, while supplementing an OP50 diet with 640nM B12 does protect against axon degeneration (N=30, 30, 8, 20). Significance relative to TIR-1(oe) on OP50 was determined by one-way ANOVA and Dunnett’s test and is indicated by ****p≤0.0001 (D, H). Significance relative to TIR-1(oe) on OP50 was determined by student’s t-test and is indicated by ****p≤0.0001 (E). Error bars represent standard deviation.

In parallel, we asked if we could leverage *C. elegans* as a model system to investigate whether individual bacteria isolated from the human microbiome can suppress Wallerian-type motor axon degeneration. We found that *Citrobacter freundii* ATCC 8090 (hereafter referred to as *Citrobacter*) colonized the *C. elegans* intestine, supported wild type growth and fecundity, and increased the number of intact motor axons in TIR-1(oe) animals (**Figure 1E**). These data demonstrate the *C. elegans* model can be used to identify neuromodulatory function of human-derived bacteria and that the *Citrobacter* diet protects motor axons from TIR-1/SARM1 induced degeneration.

Both *Comamonas aquatica* and *Citrobacter freundii* produce a significant amount of Vitamin B12 (hereafter referred to as B12). *Comamonas* species are aerobic, gram-negative, nonfermenting and mobile rod-shaped bacteria from the *Comamonadaceae* family that are abundant in the environment^14,35^. The B12 that they produce influences reproduction, developmental rate and neuronal hyperexcitability in *C. elegans*^15,36^. *Citrobacter freundii* is a facultative anaerobic, gram-negative bacteria of the *Enterobacteriaceae* family. It is a common component of the human gut microbiome that produces significant B12 such that it is used to increase B12 levels in food sources such as tempeh^37–39^. In contrast, most bacteria, including the standard lab diet *E. coli* OP50, do not produce B12. Consequently, when raised on an *E. coli OP50* diet, *C. elegans* are in a mildly B12 deficient state^40^. We hypothesized *Comamonas aquatica* and *Citrobacter freundii* may counteract TIR-1/SARM1 induced neurodegeneration by providing host animals with B12.

Although *Citrobacter freundii* is commonly found in the human microbiome, it is an opportunistic pathogen^41^. Therefore, we asked whether the nonpathogenic *Comamonas aquatica* diet protects against motor axon degeneration by providing animals with B12. We found only 2 axons were protected in animals fed a mutant *Comamonas* strain that cannot produce B12 (*cbiA*Δ) diet compared to animals fed a wild-type *Comamonas* diet, in which 9 axons were protected (**Figure 1G**). Therefore, *Comamonas* induced protection is dependent on its ability to produce B12. This finding prompted the complimentary question of whether supplementing B12 to the standard *E. coli* OP50 diet is sufficient to confer neuroprotection in TIR-1(oe) animals. It has been established that *E. coli* OP50 is able to absorb B12 from the media^15^. To determine whether TIR-1(oe) animals can absorb B12 from a supplemented OP50 diet, we visualized a previously developed B12 sensor, *Pacdh-1::GFP*. Transcription of acyl-CoA dehydrogenase (*acdh-1*) is triggered in low B12 conditions; consequently, decreased activity of the *acdh-1* promoter driving GFP expression is an accurate reflection of the presence of B12 in the animals’ diet^16^. We observed significantly less *Pacdh-1::GFP* in animals fed *E. coli* OP50 supplemented with 640 nM B12 (hereafter referred to as *E. coli* OP50 + B12) or *Comamonas* than in animals fed *E. coli* OP50, demonstrating the animals are capable of ingesting and absorbing B12 from supplemented diets (**Figure 1F**). Supplementing B12 to the *E. coli* OP50 plates was also sufficient to induce significant neuroprotection, increasing the proportion of intact motor axons from 16.3% in animals fed *E. coli* OP50 to 42.5% in animals fed *E. coli* OP50 supplemented with B12 (p≤0.0001, one-way ANOVA) (**Figure 1G, Supplementary Figure 2**). Collectively, these data demonstrate *Comamonas* protects GABAergic axons from degenerating by providing dietary B12.

### The *Comamonas* diet specifically protects against constitutively active TIR-1/SARM1

The persistent expression of active TIR-1 causes chronic insult to the nervous system, comparable to what has been found to result from constitutive SARM1 hyperactivity in chronic neurodegenerative diseases, such as ALS and Alzheimer’s^27,28^. We asked whether a *Comamonas* diet can also protect against acute insults, such as injury. To determine the scope of *Comamonas’* neuroprotective ability, we investigated established degeneration-inducing injury models in two types of neurons that have differing requirements for TIR-1/SARM1. In mechanosensory neurons, such as the PLM (posterior lateral microtubule) neuron, a single axotomy 50 µm from the cell body is sufficient to induce degeneration of the severed axon fragment. This axonal degeneration of the PLM is thought to be regulated independently of TIR-1^42^. However, a large proportion of PLM neurons are able to fuse the proximal and distal axon fragments formed by a single axotomy, thus inhibiting axon degeneration^43^. To increase the probability of injury-induced degeneration, we created a 20-40 µm gap in the PLM by severing the axon twice, which makes it more difficult for the severed axon fragments to fuse (**Supplemental Figure 3A,B**). Similarly, to investigate injury induced degeneration of GABA motor axons, we used a similar double axotomy assay, where GABA axon fragments are severed from the cell body and neuromuscular junction (**Supplemental Figure 3D)**. The majority of severed axon fragments undergo Wallerian-like degeneration that is regulated by an NADase independent interaction between TIR-1 and the DLK-1 MAP kinase pathway (**Supplemental Figure 3E)**^23,24^.

For both assays, we grew wild-type animals on *E. coli* OP50 or *Comamonas*, axotomized GFP labeled motor or mechanosensory neurons in L4 animals, and quantified the number of intact axon fragments 24 hours post injury (**Supplemental Figure 3A,D**)^44,45^. Counter to the protection observed in TIR-1(oe) animals, we found that injury-induced degeneration of mechanosensory axons occurred more often in animals raised on *Comamonas* compared to animals raised on *E. coli* OP50, indicating *Comamonas* does not protect against degeneration in this context (**Supplemental Figure 3C**). Injured GABA motor axons in wild-type animals degenerated as frequently whether they were raised on *Comamonas* or on *E. coli* OP50 (**Supplemental Figure 3E**); therefore, injury-induced degeneration of GABA axons is not prevented by a *Comamonas* diet. We also investigated whether *Comamonas* plays a role in the regenerative response after injury, which is inhibited by TIR-1. We found no change in regeneration after either single or double cut axotomy (**Supplemental Figure 3F-H**). The finding that *Comamonas* is not broadly protective across TIR-1-dependent and TIR-1-independent injury models suggests that *Comamonas* does not uniformly exert protection by downregulating TIR-1 function.

### How do *Comamonas* and B12 induced neuroprotection?

Understanding how *Comamonas* and B12 protect axons provides an opportunity to reveal molecular and cellular mechanisms that regulate axon degeneration and to identify genes that can be manipulated independently of *Comamonas* to protect against neurodegeneration. B12 is a cofactor for two proteins, methionine synthase (MET) and methylmalonyl-CoA mutase (MUT), which are highly conserved in *C. elegans* as METR-1 and MMCM-1, respectively (**Figure 2A**)^46^. METR-1 requires B12 to regulate the methionine/S-adenosylmethionine (Met/SAM) cycle, which converts homocysteine to methionine^15^. MMCM-1 requires B12 to regulate the propionate pathway to break down uneven-chain fatty acids and maintain mitochondrial homeostasis. To determine whether METR-1, MCMM-1, or both are required for the protection conferred by B12, we mutated either *metr-1(-)* or *mmcm-1(-)* in TIR-1(oe) animals and quantified the number of intact GFP labeled GABA motor axons in animals fed *E. coli OP50* or *Comamonas*. We observed significantly fewer intact axons in *metr-1(-)*;*tir-1(oe)* animals than in *tir-1(oe)* animals when either were fed *Comamonas* (p<0.0001, one way Anova with Tukey’s multiple comparisons test), (**Figure 2B**). In contrast, loss of *mmcm-1* function did not suppress protection, and loss of both *metr-1* and *mmcm-1* function in the same animal did not significantly suppress more protection than loss of *metr-1* alone, suggesting MMCM-1 does not promote B12-mediated neuroprotection. Note that in the same set of experiments, we did not observe a significant difference between the number of intact axons in *metr-1(-);tir-1(oe)* animals and *tir-1(oe)* animals fed *E. coli OP50*; therefore, *metr-1* does not influence protection in the absence of B12 (p>0.9999, one way Anova with Tukey’s multiple comparisons test). This indicates B12 requires METR-1 to protect against TIR-1(oe) induced axon degeneration.

**Figure 2:**
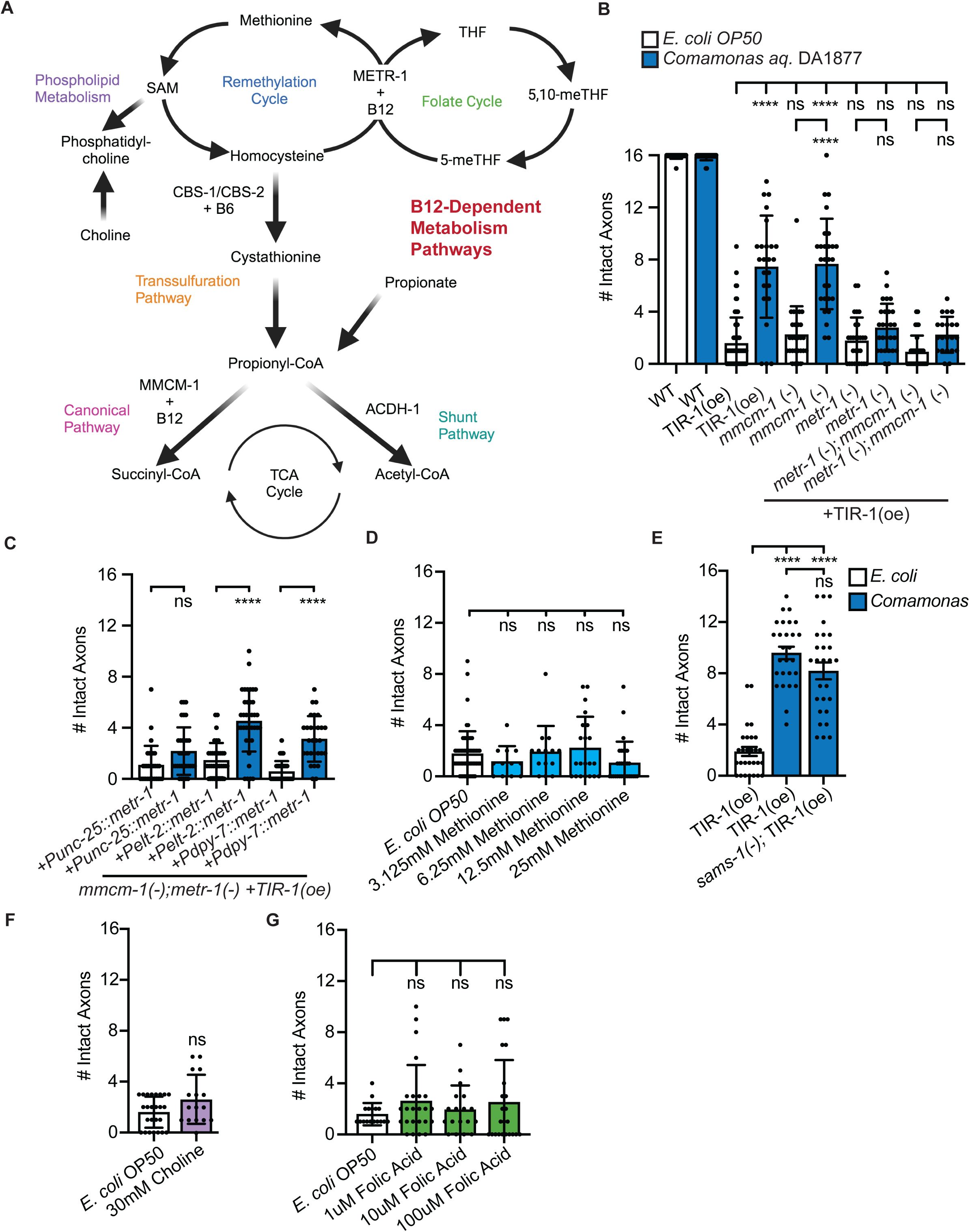
METR-1 promotes B12 mediated neuroprotection, which is regulated independently of methionine, choline or folate levels. (**A**) Overview of Vitamin B12 dependent metabolic pathways. (**B**) Neuroprotection exerted by *Comamonas* is lost in the absence of METR-1, but not MMCM-1 (N=21,23,62,24,28,27,28,27,31,21). (**C**) Cell type specific METR-1 expression in the intestine or hypodermis is sufficient to protect against TIR-1(oe) induced degeneration (N=31,33,36,37,37,29). (**D**) Adding methionine to the *E. coli* OP50 diet does not protect against TIR-1(oe) induced degeneration (N=60,12,14,21,30). (**E**) Disrupting the function of SAMS-1, a key component of phospholipid metabolism is not protective. (**F**) Supplementing the key metabolite choline in the B12 pathway does not protect against TIR-1(oe) induced degeneration (N=25,16). (**G**) Adding folic acid to the *E. coli* OP50 diet does not protect against TIR-1(oe) induced degeneration (N=17,24,18,23). Significance determined by one-way ANOVA and Tukeys’s multiple comparisons test and is indicated by ****p≤0.0001. Error bars represent standard deviation.

### Which tissues mediate *Comamonas* induced neuroprotection?

METR-1 is expressed broadly throughout the animal in the intestine, muscle, hypodermis, reproductive system, glia, and nervous system^47,48^, raising the possibility that it could function either intrinsically or extrinsically to regulate neuroprotection. We investigated where METR-1 functions to regulate B12-mediated protection by asking whether tissue-specific expression of METR-1 cDNA is sufficient to protect GABA motor axons in animals that otherwise lack METR-1 function. We expressed METR-1 driven by either GABA-, intestine-, or hypodermis-specific promoters (*Punc-25*, *Pelt-2*, *Pdpy-7*, respectively) in *metr-1(-);mmcm-1(-)* null mutants that overexpress TIR-1 (*metr-1(-);mmcm-1(-)* + *tir-1(oe)* + *Ptissuespecific:*:*metr-1*), and quantified the number of intact axons in each. Tissue specific expression in either the intestine or hypodermis was sufficient to restore axon protection to *metr-1(-);mmcm-1(-) + tir-1(oe)* animals fed *Comamonas*, while motor neuron-specific METR-1 expression did not protect axons in animals fed the same diet (**Figure 2C**). These data indicate B12 can function non-cell autonomously in peripheral tissues, such as the intestine and hypodermis, to protect TIR-1/SARM1 induced motor axon degeneration.

### B12 protects against TIR-1 induced degeneration independently of increased methionine, S-adenosylmethionine, choline, or folate

Vitamin B12 dependent METR-1 function regulates the abundance of a number of metabolites including choline, homocysteine, membrane lipids, propionate, and folic acid (**Figure 2A**). We hypothesized that supplementing metabolites that replicate the *Comamonas* phenotype on an *E. coli* OP50 diet and enzymatic mutants that block protection on the *Comamonas* diet would reveal downstream protective signaling events. Previous studies have demonstrated that proteotoxic effects of amyloid-beta and cholinergic hyperexcitability are suppressed by B12-induced increases in methionine and reduction choline levels, respectively^36,49^. We found that neither methionine nor choline supplementation suppressed axon degeneration in TIR-1(oe) animals (**Figure 2D,F**). The concentrations were chosen in accordance with those that were previously reported^48,49^. Methionine supplementation over 25 mM induced lethality in TIR-1(oe) animals. In parallel to supplementing metabolites, we mutated additional key enzymes in the MET/SAM cycle, including SAMS-1, a putative SAM synthetase gene that converts methionine to S-adenosyl methionine (SAM)^50^. Consistent with our supplementation data, we found that loss of SAMS-1 did not alter the number of degenerating axons in animals fed *Comamonas* (**Figure 2E**). In addition to being a key component of the Met/SAM cycle, METR-1 also functions in the folate metabolism cycle to convert 5-methyltetrahydrofolate (5-mTHF) to tetrahydrofolate (THF) (**Figure 2A**). Tetrahydrofolate (Vitamin B9) is the biologically active form of folate and is reduced in animals with B12 deficiency or loss of METR-1 function^15^. To determine whether *Comamonas* protects axons by providing THF, we quantified the number of intact axons in TIR-1(oe) animals fed *E. coli* OP50 that was supplemented with folic acid^15,51^, and did not observe any protection against degeneration (**Figure 2G**). The tested concentrations match those previously found to reduce egg laying and have been tested in a metabolite screen on *Pacdh-1*::GFP expression^15,51^. These data indicate B12 does not protect TIR-1(oe) axons by upregulating key metabolites of the Met/SAM cycle or through folate metabolism. Rather, together with our finding that B12 protects neurons from specific types of genetic and physical insults, these data suggest that B12 and the molecular mechanisms that function downstream of B12 are context dependent.

### *Comamonas* and B12 diets induce significant metabolic changes in TIR-1(oe) animals

Expanding our screen of candidate metabolites required a better understanding of how the metabolome of a TIR-1(oe) animal changes in response to a *Comamonas* diet. Therefore, we performed liquid chromatography-mass spectrometry (LC-MS) to identify metabolites with differential presence in animals raised on protective and non-protective diets (**Figure 3A**). To grow large age matched populations of animals with uniform TIR-1 expression, we took advantage of a strain that contains an integrated multicopy array encoding a cholinergic promoter driving the activated isoform *tir-1b* (*Punc-17*β::*tir-1b*::mCherry), *PACh::tir-1* moving forward. These *PACh::tir-1* animals express less TIR-1b in their cholinergic motor neurons than the *Pgaba::tir-1* animals and consequently, the cholinergic axons degenerate less frequently but instead exhibit the first stages of Wallerian degeneration, axon thinning and beading. In addition to being morphologically weakened, the function of the TIR-1 expressing cholinergic axons is significantly compromised compared to wild-type axons, as quantified in a thrashing assay (7.9 vs 42.3 thrashes/30 seconds, p<0.0001, student’s T-test) (**Supplementary Figure 4**).

**Figure 3:**
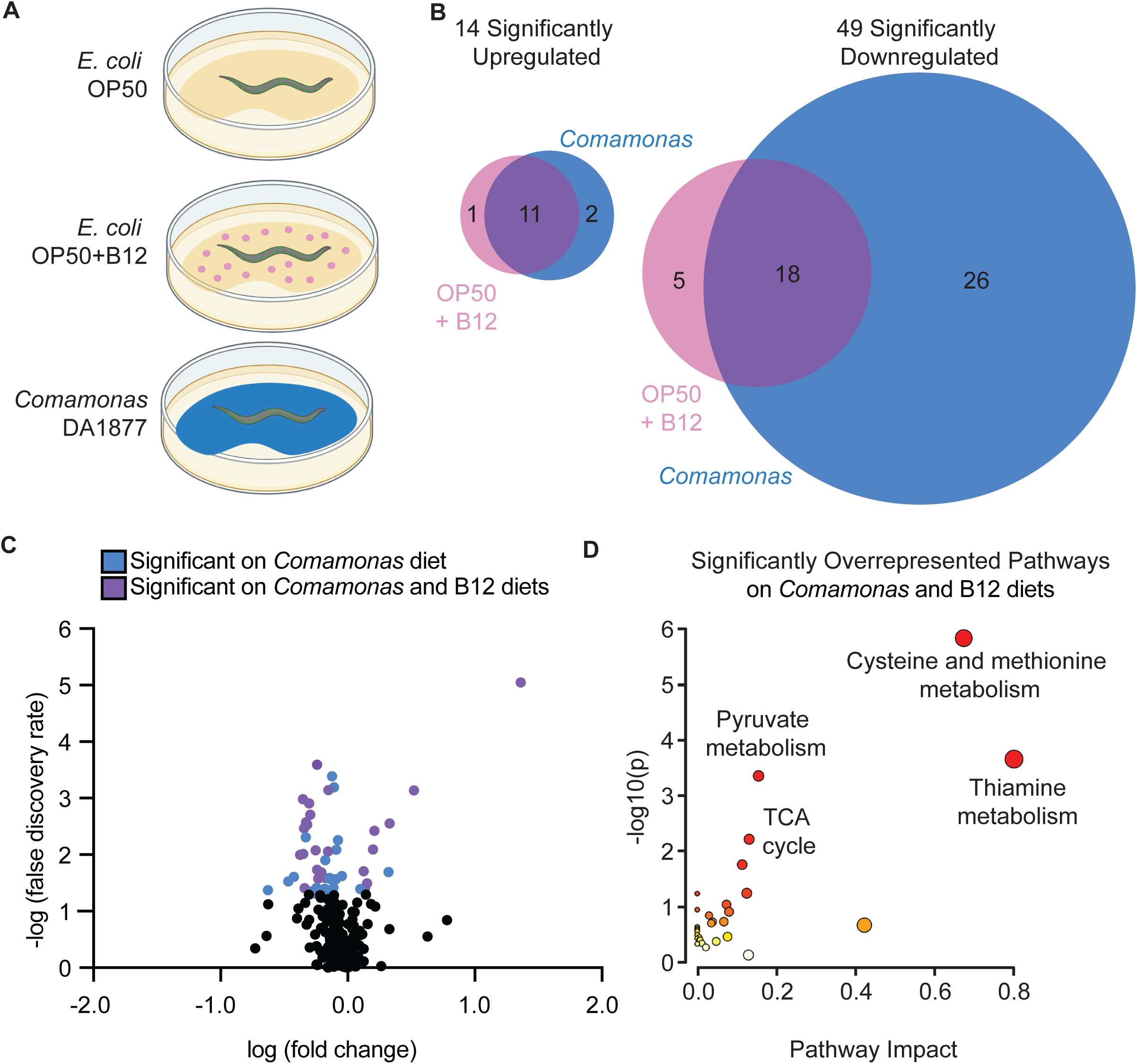
*Comamonas* and B12 diets induce significant metabolic changes in TIR-1(oe) animals. (**A**) Representative experimental set up with TIR-1 expressing animals grown up on *E. coli* OP50, *E. coli* OP50 + B12, and *Comamonas aq.* (**B**) LC-MS of whole animals grown on B12 supplemented or *Comamonas* plates demonstrates significant differences from animals grown on *E. coli* OP50 alone. 14 metabolites were upregulated on the B12 diets compared to OP50 with significant overlap. 49 metabolites were downregulated on the B12 diets compared to OP50 with significant overlap. (**C**) Polar metabolite profiling of bamIs9 [*Punc-17::tir-1b::mCherry*] animals grown on *E. coli* OP50 versus *Comamonas aq.* with significant metabolites in blue and significant metabolites on both *Comamonas aq.* and B12 supplemented plates in purple. (**D**) Metaboanalyst 6.0 pathway analysis of significantly overrepresented pathways in animals grown on a B12 diet versus *E. coli* OP50 diet with cystine and methionine metabolism as the most represented pathway. Color of circles from white to red represents degree of significance.

The LC-MS analysis of animals raised on *E. coli* OP50, *E. coli* OP50 + B12, or *Comamonas* diets detected 250 metabolites, 57 of which were significantly different in animals fed the *Comamonas* diet relative to the OP50 diet, and 35 of which were significantly different on the B12-supplemented diet relative to the OP50 diet (**Figure 3B,C**). Of the differentially represented metabolites, 29 metabolites were differentially regulated in both the OP50 + B12 and *Comamonas* datasets (**Supplemental Figure 5**). The 29 significantly altered metabolites on both *Comamonas* and B12 represent candidate regulators of TIR-1-induced degeneration. Pathway analysis of the shared metabolic profile identified multiple pathways that were significantly altered by the presence of B12 in the supplemented diet and in the *Comamonas* diet (**Figure 3D**). Not surprisingly, cysteine/methionine metabolism was the most enriched pathway as it directly utilizes B12 as a cofactor for METR-1 converting homocysteine to methionine (**Figure 2A**).

We previously found that TIR-1 induced axon degeneration relies on conserved residues required for its NADase activity, indicating that like dSARM and SARM1, activated TIR-1 induces axon degeneration by hydrolyzing NAD^+23,24,27,52–55^. Prominent biproducts of TIR-1 mediated NAD^+^ hydrolysis are nicotinamide (NAM), adenosine diphosphate ribose (ADPR), and cyclic adenosine diphosphate ribose (cADPR) (**Supplementary Figure 6A**)^52,55^. Therefore, we asked whether *Comamonas* suppresses or counteracts the pro-degenerative NADase function of TIR-1 by comparing the amount of available NAD^+^, NAM, ADPR and cADPR in *PACh::tir-1* animals fed *E. coli* OP50*, E. coli* OP50 *+ B12, or Comamonas.* We found that NAD^+^, cADPR, ADPR, and NAM levels are not significantly downregulated in animals fed either B12 containing diets relative to *E. coli* OP50, although there is a trend towards a reduction in all four metabolites, (**Supplementary Figure 6B-E**). Although the diets were processed in parallel, it appears that the lack of significance is due to variability in metabolite levels in animals raised on *E. coli* OP50. Each of the NAD^+^ byproducts are more tightly regulated in the presence of either B12-containing diet, indicating there may be a small interaction between expression of TIR-1 and a B12 diet; however, the nature of that relationship remains unknown. Nonetheless, the lack of significance in our comparison of NAD^+^ byproducts suggests B12 does not solely regulate degeneration by reducing the NADase function of TIR-1.

### B12 exerts neuroprotection by lowering toxic homocysteine levels

From our metabolomic analysis we also found that methionine and S-adenosylmethionine were significantly increased in animals grown on both B12 and *Comamonas* diets, while homocysteine was significantly decreased (**Figure 4A-C**). These results are consistent with previous gas chromatography-mass spectrometry data in wild-type animals^56^. Homocysteine is toxic to neurons and its accumulation is a key biomarker of Alzheimer’s disease^57,58^. The significant decrease in homocysteine generated the hypothesis that *Comamonas* may protect against degeneration by decreasing homocysteine levels in response to B12. Therefore, we asked whether lowering homocysteine levels independently of B12 would also inhibit TIR-1(oe) induced degeneration. In addition to being used as a substrate for methionine synthesis by B12 and METR-1, homocysteine is converted to cystathionine in the presence of Vitamin B6 in the transsulfuration pathway (**Figures 2A, 4D**)^59^. We found that supplementing B6 to TIR-1(oe) animals led to a small but significant increase in axon protection relative to TIR-1(oe) animals raised on an *E. coli* OP50 diet (**Figure 4E**). This indicates that lowering homocysteine levels is sufficient to protect axons, even in the absence of B12. To further test our hypothesis, we asked whether supplementing homocysteine to plates seeded with *E. coli* OP50 would exacerbate axon degeneration. This experiment could not be done with the TIR-1(oe) animals, since axon degeneration on an *E. coli* OP50 diet was virtually complete. Instead, we asked whether supplementing *E. coli* OP50 with homocysteine would induce degeneration in wild-type animals or in low expressing pGABA::TIR-1 animals that express relatively low amounts of activated TIR-1. We found that homocysteine did not induce degeneration in wild-type L4 animals or in wild-type adult animals aged 3 days beyond L4. However, a significant number of pGABA::TIR-1 day 3 adults had broken axons and gaps in their dorsal nerve cord, which were significantly more prevalent in the presence of homocysteine (**Figure 4F,G**). These data demonstrate that homocysteine is specifically toxic to aged TIR-1(oe) neurons, and that by supplying B12 and decreasing homocysteine levels, *Comamonas* promotes neuroprotection in the presence of constitutive TIR-1/SARM1 NADase function (**Figure 4H**).

**Figure 4:**
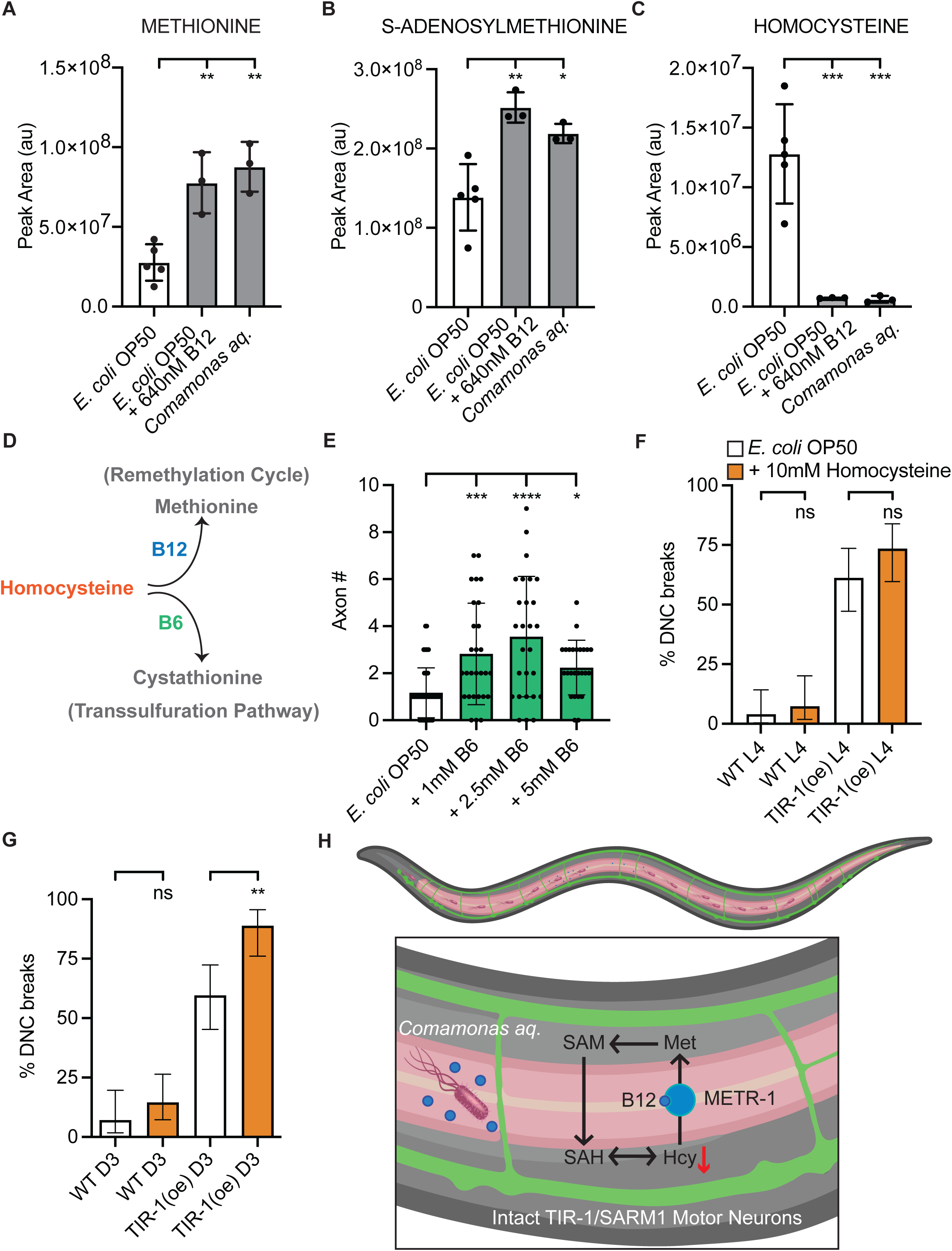
*Comamonas* and B12 protect by lowering toxic homocysteine levels. **(A)** LC-MS data from animals fed *E. coli* OP50 versus *E. coli* OP50 + B12 or *Comamonas* demonstrates a significant increase in methionine in animals grown in the presence of B12. **(B)** LC-MS data from animals fed *E. coli* OP50 versus *E. coli* OP50 + B12 or *Comamonas* demonstrates a significant increase in S-adenosylmethionine in animals grown in the presence of B12. **(C)** LC-MS data from animals fed *E. coli* OP50 versus *E. coli* OP50 + B12 or *Comamonas* demonstrates a significant decrease in homocysteine in animals grown in the presence of B12. Significance relative to wild type was determined by one-way ANOVA and Dunnett’s test and is indicated by *≤0.05, **p≤0.01, and ***p≤0.001. Error bars represent standard deviation. (**D**) Homocysteine feeds into both the remethylation cycle and the transsulfuration pathway, depending on the presence of Vitamins B12 and B6, respectively. (**E**) Supplementing Vitamin B6 to *E. coli* OP50 protects against TIR-1(oe)-induced axon degeneration (N=48,28,27,25). Significance was determined by one-way ANOVA and Dunnett’s test and is indicated by ****p≤0.0001, ***p≤0.001, *p≤0.05. Error bars represent standard deviation. (**F**) Supplementing homocysteine to animals that express sub threshold amounts of TIR-1 does not change the proportion of L4 animals with broken axons and gaps in their dorsal nerve cords (N=50,41,49,49). Significance was determined by Fischer’s exact test. Error bars represent 95% confidence intervals. (**G**) Supplementing homocysteine to the same TIR-1 expressing animals specifically increases the proportion of Day 3 adult animals with axon breaks and gaps in their dorsal nerve cords (N=42,55,47,45). Significance was determined by Fischer’s exact test and is indicated by **p≤0.01. Error bars represent 95% confidence intervals. (**H**) Model of B12 mediated neuroprotection. *Comamonas* produces B12, which improves neuronal integrity though METR-1, by lowering toxic homocysteine levels.

## Discussion

The microbiome has recently been identified as a promising target in modulating nervous system function and protection. However, which gut bacteria and their metabolites exert protection and the molecular mechanisms by which they impact the nervous system remain largely unknown. The well characterized nervous system and the ability to exist on a uni-bacterial diet makes *Caenorhabditis elegans* one of very few models in which interactions between specific bacteria and the nervous system can be explored in a disambiguated manner. In this study we found *Comamonas aquatica* DA1877 protects against motor axon degeneration that is induced by chronic expression of a key and conserved prodegenerative gene, TIR-1/SARM1. We found that *Comamonas’* neuroprotective ability relies on its ability to produce Vitamin B12, and on B12-dependent METR-1/MET methionine synthase function in the intestine, to lower toxic homocysteine levels. In addition, we developed a novel protocol to identify single microbiota isolated from human intestines that suppress axon degeneration.

### Overexpression of activated TIR-1 is a tractable model to identify gut-brain interactions

Screening uni-bacterial diets with a *C. elegans* model of degeneration enabled us to uncover multiple diet derived aerobic and anaerobic neuroprotective bacteria, identify bacterially provided neuroprotective metabolites, as well as identify responding genes and metabolites that function within the host to confer neuroprotection. TIR-1 is highly conserved with mammalian SARM1 and degeneration in TIR-1(oe) animals requires the conserved NADase mechanism of action of its SARM1 orthologs^23,24,53^. Previous studies have identified SARM1 variants that constitutively hyperactivate its NADase function in patients with neurodegenerative disease^25–31^. Therefore, novel strategies to inhibit constitutive SARM1 expression and activity are highly sought after as potential therapeutic interventions. Here, by overexpressing a short isoform of TIR-1 that lacks its N-terminal autoinhibitory domain, the NADase activity of TIR-1 is also constitutively activated. In light of the finding that *Comamonas* and B12 diets did not significantly affect NAD^+^ hydrolysis, the data demonstrate that the gut-brain axis can be manipulated to modulate TIR-1 induced degeneration by intervening downstream or in parallel to TIR-1 activity and function. However, TIR-1 overexpression is confined to the cholinergic neurons, so metabolic changes in the neurons may be obscured by whole animal analyzes.

Expanding the screen to include bacteria isolated from the human intestine revealed a new method to identify neuroprotective bacteria. Determining the mechanism of *Citrobacter freundii*-induced axon protection will be an intriguing avenue of future investigation. We hypothesize *Citrobacter freundii* also protects axons by supplying B12 and lowering homocysteine levels. However, it is also possible that *Citrobacter freundii* protects axons by producing exopolysaccharides, which are powerful antioxidants. Our data demonstrate that we can 1) reconstitute complex human microbiomes in the animal; 2) identify individual human microbiota that can colonize the animal intestine and support *C. elegans* growth; and 3) identify human microbiota that protect against neurodegeneration. Therefore, we have established crucial proof-of-principle that diverse human microbiota can colonize the *C. elegans* intestine and that a humanized *C. elegans* microbiome can be used to investigate mechanisms of neuronal function and neurodegeneration.

Together, the results demonstrate the TIR-1(oe) animals are an extremely powerful invertebrate model for in-depth molecular dissection of microbiome-host interactions. Identifying additional microbiota that can colonize the *C. elegans* intestine and influence the progression of degeneration in tractable models of neurodegenerative disease will reveal additional B12- and homocysteine-independent regulators of axon degeneration. Given the complex interactions between bacteria within the microbiome, screening defined combinations of bacterial diets will also be highly informative.

### *Comamonas aquatica* protects against specific types of neuronal insult

Our understanding of canonical SARM1 function is that it is activated after acute injury to promote degeneration of severed axon fragments. While *Comamonas* exerts strong protection of TIR-1(oe) axons, we did not observe changes in protection or repair after acute injury to GABA motor neurons (**Supplementary Figure 1**). Moreover, the enhanced degeneration observed in injured mechanosensory neurons in animals fed a *Comamonas* diet highlights that not only is the protective effect of *Comamonas* specific to the constitutively active model of degeneration, the *Comamonas* diet can both inhibit and promote injury induced degeneration in specific types of neurons. Understanding the context specific molecular mechanisms of bacterial supplied neuroprotection will be critical for designing effective therapies.

We found that *Comamonas* and OP50 + B12 diets lower homocysteine levels, that supplementing vitamin B6, which also lowers homocysteine levels, protects axons, and that supplementing homocysteine specifically enhanced degeneration in the presence of constitutively activated TIR-1 ^60,61^. Our data suggest animals raised on a regular *E. coli* OP50 diet are living in a state with elevated homocysteine levels, similar to the condition Hyperhomocysteinemia, which is a condition with wide ranging and debilitating consequences to human physiology, including cardiovascular and neurodegenerative disease^62^. While the observed levels of homocysteine in wild type animals fed *E. coli* OP50 are not sufficient to induce degeneration on their own, they appear to sensitize animals to a neurodegenerative trigger such as constitutive activity of TIR-1. Our data also agree with an existing finding that supplementation of B6 vitamins slows brain atrophy in elderly people with mild cognitive impairment^63^.

In parallel, low levels of B12 can lead to Subacute combined degeneration and dementia, which can be reversed upon restoration of healthy B12 levels^64,65^. Although a negative correlation between B12 levels and the presence of neurodegenerative disease exists, simply providing B12 to patients has largely failed to protect the nervous system in clinical studies^66,67^. We found that there is a specific interaction between B12, homocysteine and constitutively active TIR-1 in the aged nervous system. In contrast, hyperexcitability and proteotoxicity are reduced by B12-dependent changes in choline and methionine levels, respectively^36,49^. Surprisingly, removal of dietary B12 or addition of propionate can also suppress neurodegeneration of dopaminergic neurons in an alpha-synuclein model of degeneration in *C. elegans*^68^. These data demonstrate the context specific nature and complexity of B12’s role in neuroprotection and reflect the importance of identifying specific molecular interactions between the microbiome, its metabolites, host genes and metabolites with specific neuronal insults.

### Additional mechanisms of *Comamonas aquatica* induced neuroprotection

In addition to homocysteine, our LC-MS data identify additional metabolites that are differentially regulated in animals fed *Comamonas* or *E. coli* OP50 + B12 diets relative to *E. coli* OP50. Because B12 supplementation does not fully recapitulate the amount of protection conferred by a *Comamonas* diet, it is likely that one or some of these metabolites also contribute to neuroprotection. Similarly, B6 supplementation conferred less protection than B12, suggesting the amount of B6 provided to the intestine may inefficiently reduce homocysteine levels or that B12 protects against degeneration using one or more mechanisms in addition to reducing homocysteine.

Taken together, we have identified bacterial diets that strongly protect against TIR-1/SARM1 induced neurodegeneration. *Comamonas* had the strongest ability to protect GABA motor axons from degenerating, and this protection is mediated by Vitamin B12 and the enzyme METR-1, which decrease neurotoxic homocysteine. This study highlights *C. elegans* as a model to not only identify specific neuroprotective bacteria, but also elucidate complex and conserved molecular mechanisms of neuroprotection within the bacteria and host.

## Acknowledgments

We thank the M Francis and E Yemini labs, M Walhout, B McCormick, and members of the Byrne lab for helpful discussions. We also thank P Emery and D Winder for helpful comments on the manuscript. This work was supported by grants from the Riccio Fund for Neuroscience and the National Institutes of Health (NINDS F31 NS124338-01A1 and NINDS R01 NS110936-01) to L.C.O. and A.B.B.. Some bacterial strains used in this work were provided by the CGC.

## Author contributions

Conceptualization, L.C.O. and A.B.B.; formal analysis, L.C.O.; investigation, L.C.O. and W.K.K.; methodology, L.C.O., W.K.K., and P.V.; visualization, L.C.O.; funding acquisition, L.C.O. and A.B.B.; product administration, A.B.B.; resources, A.B.B., M.J.A., and J.B.S.; supervision, A.B.B.; writing-original draft, L.C.O. and A.B.B.; writing-review & editing, L.C.O., A.B.B., W.K.K., P.V., M.J.A., and J.B.S.

## Figure Legends

**Supplementary Figure 1:**
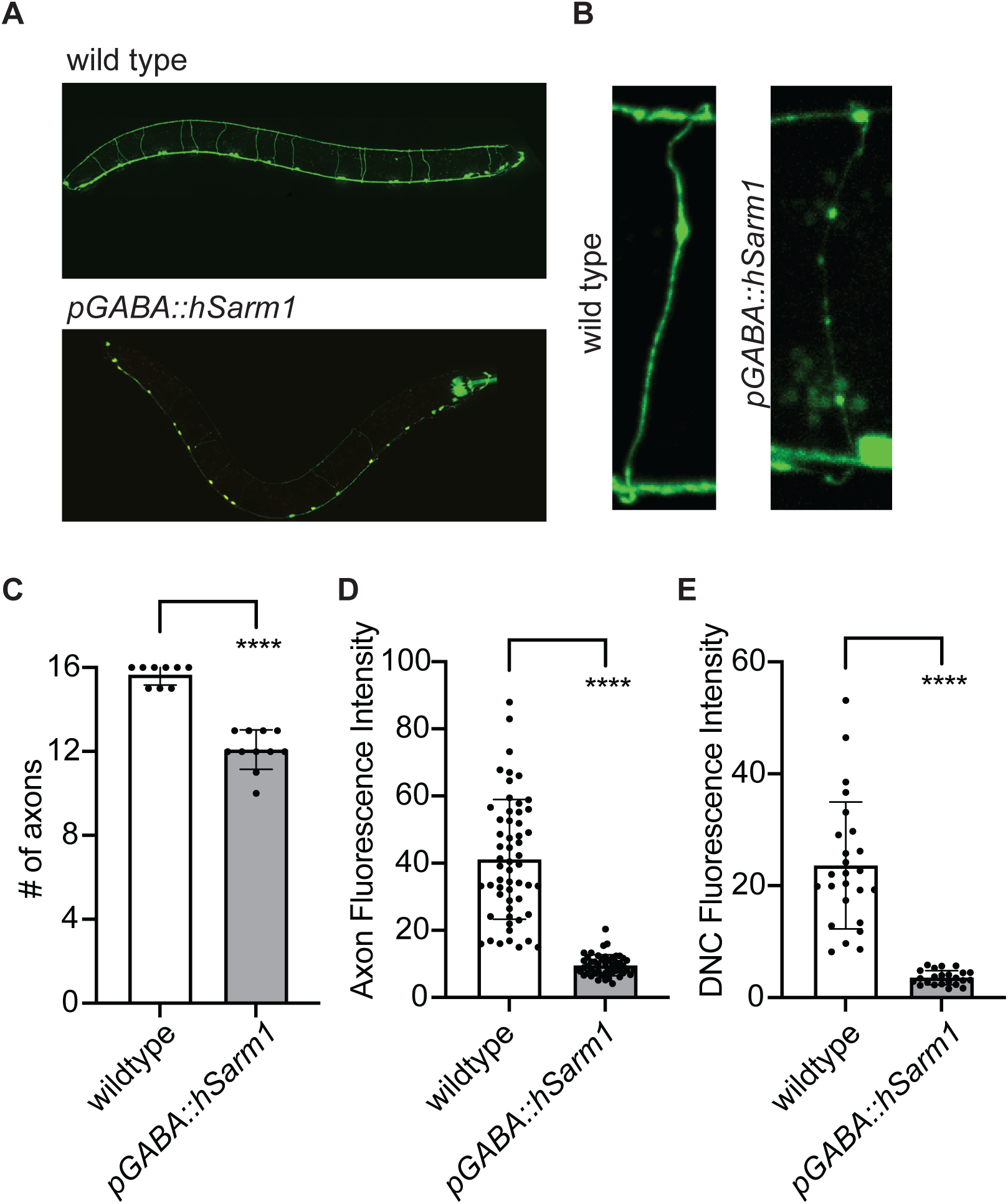
Constitutive expression of activated human SARM1 in *C. elegans* motor neurons induces significant degeneration. (**A**) Representative image of degeneration induced by hSARM1 expression in GABA motor neurons. (**B**) Representative image of beading axon in hSARM1 animals and control intact axon. The number of intact axons (**C**), fluorescence intensity of motor axon commissures (**D**), and fluorescence intensity of axons in the dorsal nerve cord (**E**). Significance relative to wildtype was determined by student’s t-test and is indicated by ****p≤0.0001. Error bars represent standard deviation.

**Supplementary Figure 2:**
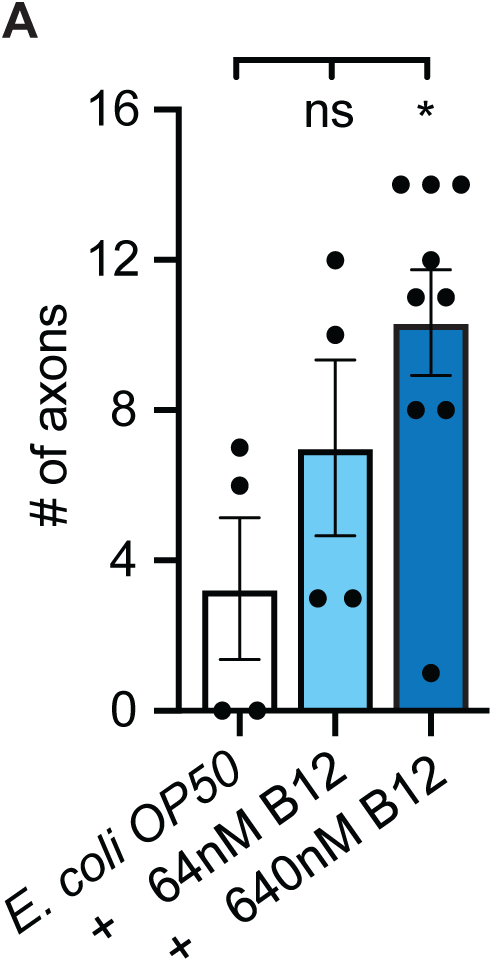
B12-induced neuroprotection is concentration dependent. (**A**) B12 supplementation protects against TIR-1 mediated degeneration in a dose dependent manner (N=4,4,9). Significance relative to wild type was determined by one-way ANOVA and Dunnett’s test and is indicated by *p≤0.05. Error bars represent standard error of the mean.

**Supplementary Figure 3:**
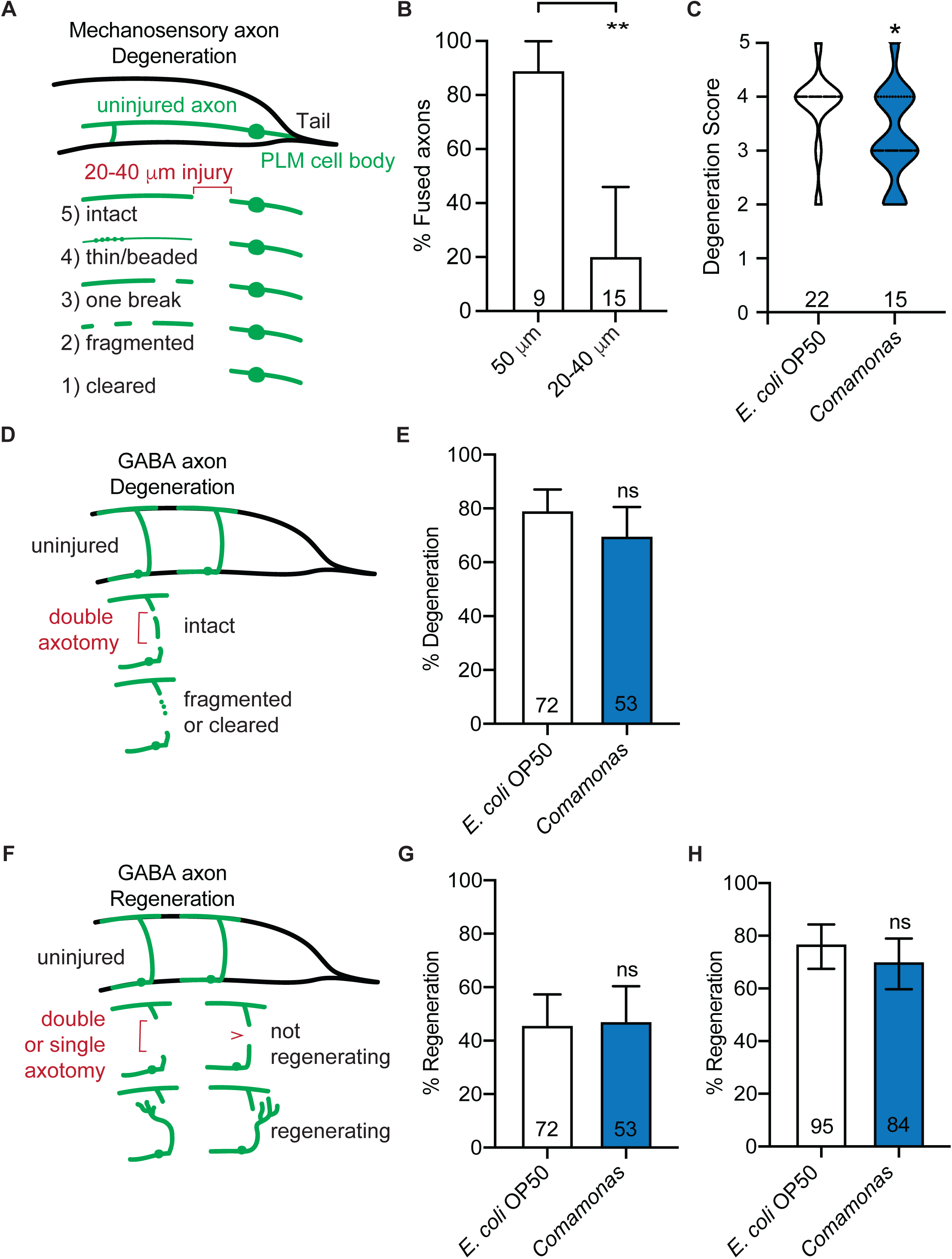
B12 diet is not broadly protective against acute injury. (**A**) Schematic representation of axotomy of PLM mechanosensory neuron and scoring scheme. (**B**) Significantly fewer PLM axons fuse after double cut axotomy (N=9,15). (**C**) PLM axons degenerate significantly more when fed a *Comamonas* diet. (**D**) Schematic representation of double cut axotomy and degeneration of GABA motor neurons and scoring scheme. (**E**) Degeneration of the middle fragment is not altered by a B12 diet (N=72,53). (**F**) Schematic representation of single and double cut axotomy for regeneration of GABA motor neurons and scoring scheme. (**G**) Regeneration after double cut injury is not altered by a B12 diet (N=72,53). (**H**) Regeneration after single cut injury is not altered by a B12 diet (N=95,84). Significance was determined by Fischer’s exact test and is indicated by *p≤0.05. Error bars represent 95% confidence intervals.

**Supplementary Figure 4:**
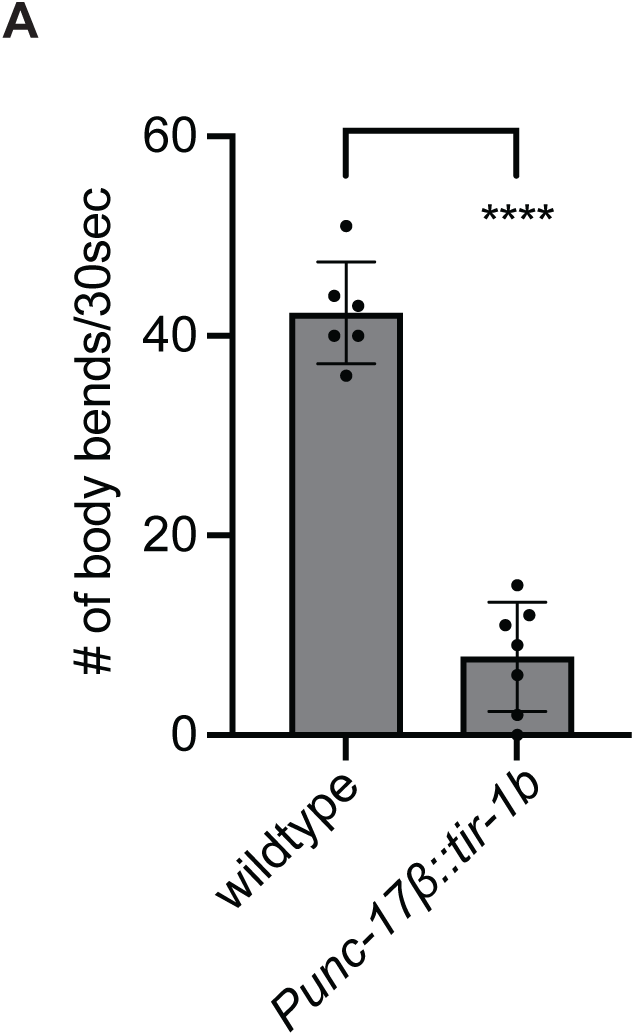
Low expression of TIR-1 in cholinergic neurons impairs neuronal function. (**A**) Integrated TIR-1(oe) in cholinergic neurons (bamIs9) decreases animals’ ability to thrash (N=6,7). Significance was determined by student’s t-test and is indicated by ****p≤0.0001. Error bars represent standard deviation.

**Supplementary Figure 5:**
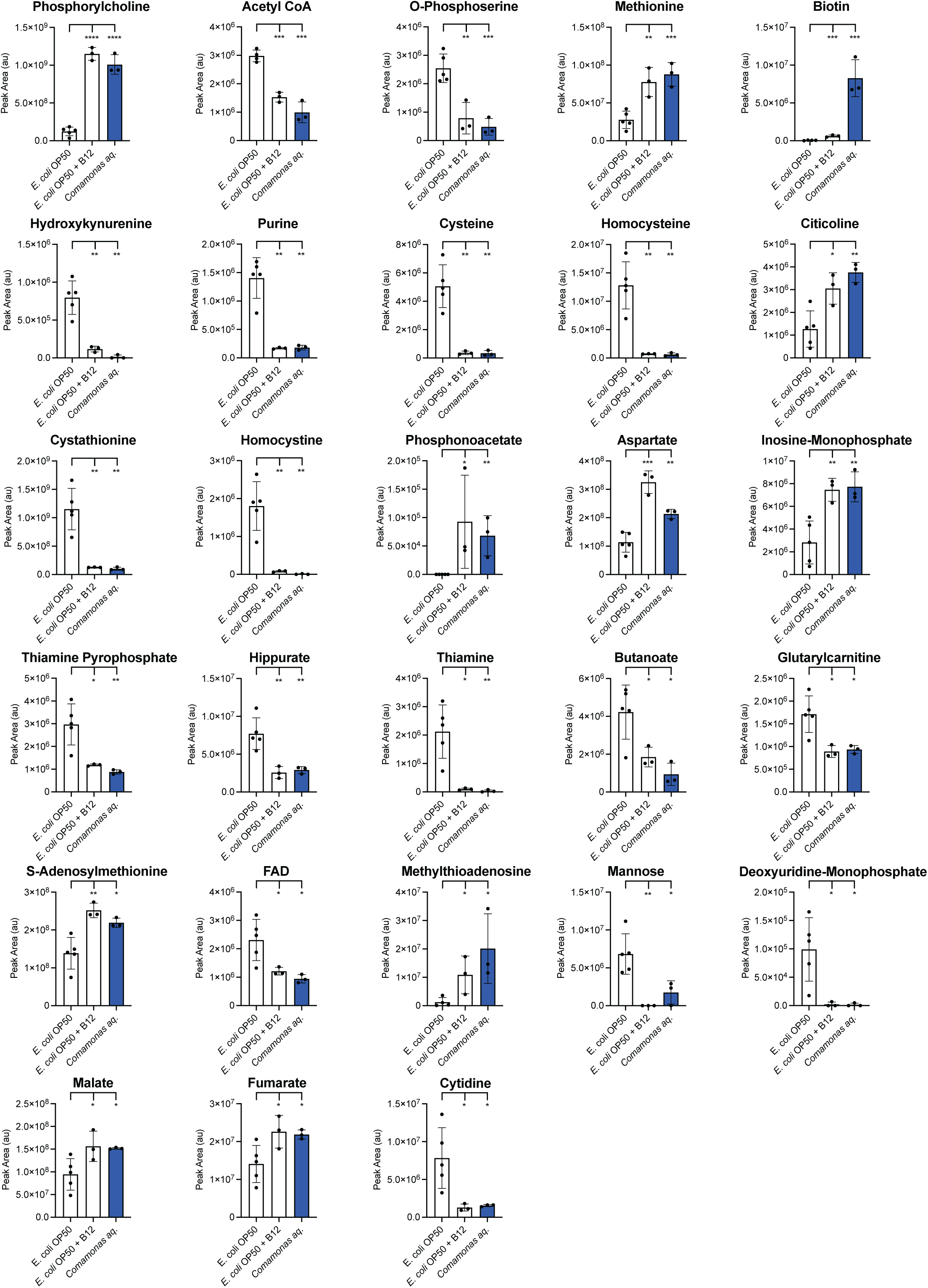
A B12 diet induces metabolic changes in *C. elegans.* The relative abundance of 29 metabolites were significantly altered on B12 diets.

**Supplementary Figure 6:**
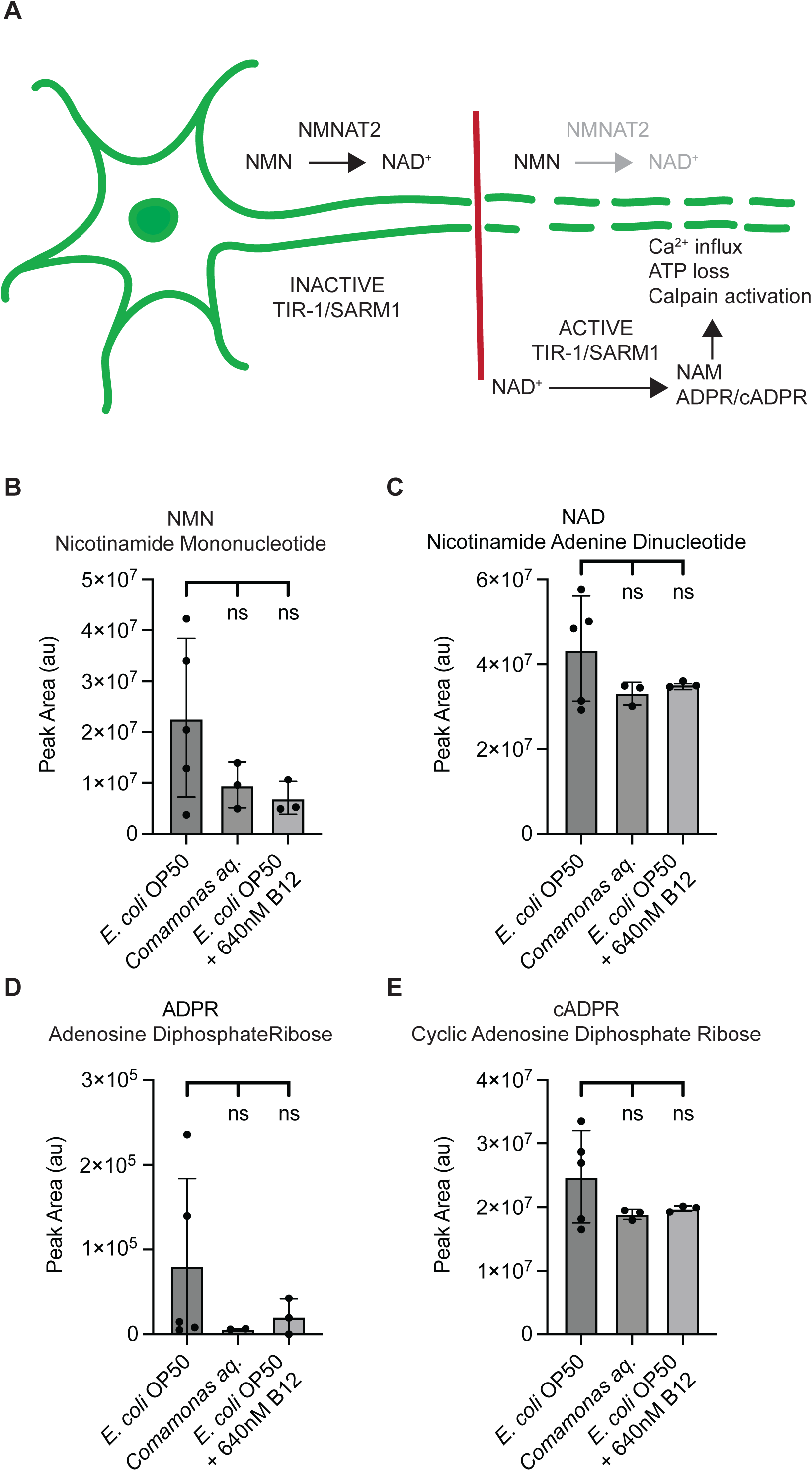
B12 does not significantly alter TIR-1 metabolites. (**A**) Simplified diagram of TIR-1/SARM1 activity. (**B**) Nicotinamide mononucleotide, a key metabolite upstream of TIR-1 activation is not significantly changed on B12 diets (N=5,3,3). (**C**) Nicotinamide adenine dinucleotide, a key metabolite TIR-1 hydrolyzes upon activation is not significantly changed on B12 diets (N=5,3,3). (**D**) Adenosine diphosphate riboside, a key metabolite produced upon TIR-1 activation is not significantly changed on B12 diets (N=5,2,3). (**E**) Cyclic adenosine diphosphate riboside, a key metabolite produced upon TIR-1 activation is not significantly changed on B12 diets (N=5,3,3). Significance relative to wild type was determined by one-way ANOVA and Dunnett’s test. Error bars represent standard deviation.

## Methods

### Nematode strains

*C. elegans* strains were obtained from the *Caenorhabditis* Genetics Center or generated in this study. All strains were maintained at 20°C on nematode growth medium (NGM) plates seeded with specific bacterial diets. Transgenic strains were generated via standard microinjection of the transgene and a fluorescent co-injection marker.

### Bacterial culture

Liquid cultures of the standard laboratory diet *Eschericia coli* OP50, *Comamonas aquatica* DA1877, and *Citrobacter freundii* ATC 8090 strains were grown at 37°C in Luria Broth overnight, while *Raoultella* sp. Jub54 and *Providencia* sp. Jub39 were grown at 30°C overnight. *Raoultella* sp. Jub54 was previously identified as *Providencia* sp. JUb39^13,69^. All were seeded onto NGM plates and left for 48 hours before transferring animals to the bacterial lawns.

### *Citrobacter freundii* isolation and identification

To isolate human fecal bacteria that colonize the gut of *C. elegans*, freshly collected feces were suspended in 50 ml of PBS per gram of feces. Solid particles were pelleted by centrifugation, and 100 μl of the supernatant was plated onto NGM plates. L4 stage animals were then transferred onto the plates and allowed to grow at 20 °C for 24hr. Approximately 30 young adult animals were picked for gut bacteria isolation, incubated in a 1/50 dilution of bleaching solution at room temperature for 10 min to remove bacteria from worm surface, washed four times with 1 ml cold M9, and lysed using 0.5 mm glass beads in a bead-beater for 3 minutes at maximum speed. The lysates were diluted in M9 and plated on LB agar plates, where different bacterial colonies were isolated based on color, shape, and size after overnight incubation at 37°C. The bacterial colonies were identified by sequencing the 16S rRNA gene using 27f (AGAGTTTGATCMTGGCTCAG) and 1492r (AAGTCGTAACAAGGTAACC) primers.

### Metabolite supplementation

Stock solutions of B12 (3.2 mM), methionine (250 mM), choline (1 M), homocysteine (500 mM), and B6 (Pyridoxal-5-phosphate) (10mM) were diluted in distilled H_2_O, while folic acid (100mM) was diluted in 1M NaOH. For each experiment, stock solutions were diluted to the desired concentration in NGM agar before pouring the plates. Methionine supplementation over 25 mM induced lethality in TIR-1(oe) animals.

### Degeneration

Chronic degeneration was assessed by immobilizing L4 stage animals with 300mM sodium azide on a 3% agarose pad. Motor axon integrity was qualitatively scored as intact, thinning/beading, broken, or absent. Intact and thinning/beading axons were combined to form the connected axons group, these axons all reach the dorsal nerve cord. Images were taken on a Perkin Elmer Precisely UltraVIEW VoX spinning disc confocal imaging system using a 40x objective.

### Laser axotomy

GABAergic motor neurons were axotomized as previously described using a pulsed nitrogen laser ^23,44^. Axon degeneration of the severed middle fragment was quantified 24hr after injury by placing animals immobilized with 300 mM sodium azide on a 3% agarose pad. Mechanosensory PLM axons were severed approximately 20 and 40μm anterior to the cell body to avoid cell-cell fusion, and the integrity of the severed distal fragment was scored 72hrs post injury. Degeneration of mechanosensory neurites was quantified according to previously described criteria ^42^. Animals were imaged with a Nikon 100x 1.4 NA objective, Andor Zyla sCMOS camera and Leica EL6000 light source.

### Molecular Biology

To determine which tissue METR-1 is sufficient to rescue B12 mediated protection, we built transgenic animals expressing METR-1 from the *unc-25* GABA-specific promoter, *dpy-7* hypodermal promoter, and *elt-2* intestinal promoter. The starting METR-1 plasmids (*metr-1p*::METR-1::*unc-54* 3’UTR, *elt-2p*::METR-1::*unc-54* 3’UTR, and *dpy-7p*::METR-1::*unc-54* 3’UTR) were a kind gift from the Alkema lab at UMass Medical School. The *dpy-7* promoter of the *dpy-7p*::METR-1::*unc-54* 3’UTR plasmid was cut out and the *unc-25* promoter from PLZ032 (*unc-25p*::mCherry) was cloned using Gibson cloning. The resulting *unc-25p*::METR-1::*unc-54* 3’UTR plasmid was injected into ABC490(*metr-1(ok521)*; *mmcm-1(ok1631)*; Ex[unc-47p::TIR-1b::*unc-54* 3’UTR]; oxIs12(*unc-47p*::GFP) at a final concentration of 10ng/µL to generate bamEx5000. The other strains were made in the same fashion injecting 10ng/µL of *elt-2p*::METR-1::*unc-54* 3’UTR (bamEx5001) or *dpy-7p*::METR-1::*unc-54* 3’UTR (bamEx5004) into ABC490.

### Imaging and fluorescence quantification

To assess neuronal integrity animals were mounted on 3% agarose pads and immobilized with 300mM sodium azide. Images were taken on a Perkin Elmer Precisely UltraVIEW VoX spinning disc confocal imaging system using a 40x objective. Images were stitched together, and axon integrity was scored in a binary way as intact or broken. Dorsal nerve cord integrity was analyzed by tracing the dorsal nerve cord using a segmented line ROI and quantifying the mean intensity.

### Liquid Chromatography-Mass Spectrometry

Approximately 3,000 animals were collected with ice cold M9 buffer from large plates and centrifuged at 1300rcf for 3 min. Supernatant was carefully removed and animals were washed 3 times more with 10ml cold M9 each time. Animals were transferred with 1ml cold M9 to 1.7ml Eppendorf tube and spun down (1300rcf, 3min). We removed as much supernatant as possible and transferred the ∼3,000 animals into 2 mL FastPrep Tubes with ceramic beads. Animals were sonicated in 1 mL precooled 80% Methanol 20% water (LCMS Grade) for 2 minutes. Then animals were pulverized in FastPrep machine 3 times 1 minute, resting on ice for 5 minutes between each pulse. Samples were vortexed at 4°C for 10 min and centrifuged at top speed for 10 min at 4°C. Supernatant was transferred to a new tube, and samples were dried using a SpeedVac (metabolites sample). Protein pellets were resuspended in 1mL RIPA buffer and vortexed at 4°C for 10 min and centrifuged at top speed for 10 min at 4°C. BSA assay was performed on protein to determine amount of protein in each sample. Dried metabolite pellets were resuspended in water to a concentration of 1 µg/µL (based on BSA), vortexed for 10 min at 4 °C and spun for 10 min at top speed at 4 °C, and supernatant from each sample was transferred to a liquid chromatography−mass spectrometry (LC–MS) vial (Thermo Fisher Scientific, 6ESV9-04PP and 6ASC9ST1). Data were collected using a Q Exactive Plus Orbitrap mass spectrometer equipped with a HESI II probe and quantified by integrating peaks in TraceFinder 5.1 (Thermo Fisher Scientific). Mass tolerance was set to 5 ppm, and expected retention times were benchmarked using in-house chemical standards. Significantly altered metabolites were analyzed using Metaboanalyst 6.0 to identify overrepresented pathways^70^.

